# Mapping Stress-Responsive Signaling Pathways Induced by Mitochondrial Proteostasis Perturbations

**DOI:** 10.1101/2024.01.30.577830

**Authors:** Nicole Madrazo, Zinia Khattar, Evan T. Powers, Jessica D. Rosarda, R. Luke Wiseman

## Abstract

Imbalances in mitochondrial proteostasis are associated with pathologic mitochondrial dysfunction implicated in etiologically-diverse diseases. This has led to considerable interest in defining the biological mechanisms responsible for regulating mitochondria in response to mitochondrial stress. Numerous stress responsive signaling pathways have been suggested to regulate mitochondria in response to proteotoxic stress, including the integrated stress response (ISR), the heat shock response (HSR), and the oxidative stress response (OSR). Here, we define the specific stress signaling pathways activated in response to mitochondrial proteostasis stress by monitoring the expression of sets of genes regulated downstream of each of these signaling pathways in published Perturb-seq datasets from K562 cells CRISPRi-depleted of individual mitochondrial proteostasis factors. Interestingly, we find that the ISR is preferentially activated in response to mitochondrial proteostasis stress, with no other pathway showing significant activation. Further expanding this study, we show that broad depletion of mitochondria-localized proteins similarly shows preferential activation of the ISR relative to other stress-responsive signaling pathways. These results both establish our gene set profiling approach as a viable strategy to probe stress responsive signaling pathways induced by perturbations to specific organelles and identify the ISR as the predominant stress-responsive signaling pathway activated in response to mitochondrial proteostasis disruption.

## INTRODUCTION

The regulation of mitochondrial protein homeostasis (proteostasis) is critical to maintain the activities of essential mitochondrial functions including energy production, lipid metabolism, and the regulation of apoptotic signaling. Mitochondrial proteostasis is maintained by a network of mitochondrial-localized protein import factors, chaperones/folding enzymes, and proteases that coordinate to facilitate the targeting, folding, and degradation of the ∼1100 proteins localized to this organelle.^1-4^ Imbalances in mitochondrial proteostasis induced by genetic, environmental, or aging-related factors promote pathologic mitochondrial dysfunction implicated in the onset and pathogenesis of etiologically diverse diseases including cancer, metabolic disease, and neurodegenerative disease.^4-9^ This has led to significant interest in understanding the molecular mechanisms involved in regulating and adapting mitochondrial proteostasis in response to pathologic insults.

Numerous stress-responsive signaling pathways have been suggested to be activated in response to cellular stress and regulate mitochondrial proteostasis. Of these, the mitochondrial unfolded protein response (UPR^mt^) has received the most attention. While originally identified as a distinct stress-responsive signaling pathway in *C. elegans*^10^, in mammalian cells, the UPR^mt^ signals through the broader integrated stress response (ISR).^11,12^ The ISR comprises four stress-responsive eIF2α kinases – PKR, GCN2, HRI, and PERK – that are activated in response to diverse cellular insults including viral infection, amino acid deprivation, heme deficiency, and endoplasmic reticulum (ER) stress, respectively.^13,14^ Once activated, these ISR kinases signal through a mechanism involving selective phosphorylation of eIF2α, leading to both a transient attenuation in new protein synthesis and the activation of stress-responsive transcription factors such as ATF4.

Previous results show that the ISR regulates many aspects of mitochondrial proteostasis downstream of eIF2α phosphorylation. ISR-dependent activation of ATF4 induces expression of multiple mitochondrial proteostasis factors including the HSP70 chaperone *HSPA9* and the AAA+ quality control protease *LON*.^15-18^ Further, ISR-dependent translational attenuation regulates mitochondrial protein import through the degradation of the core import subunit TIM17A.^18,19^ ISR signaling has also been implicated in the regulation of many other aspects of mitochondrial biology including energy metabolism, lipid regulation, and organellar morphology.^18,20-24^ These results highlight a central role for the ISR in regulating mitochondria in response to stress.

Recent work has revealed the molecular mechanism by which mitochondrial stress activates ISR signaling. In response to mitochondrial stress, the stress-sensing mitochondrial protein DELE1 accumulates in the cytosol as either a full-length protein or proteolytic fragment generated by the stress-activated inner membrane protease OMA1.^25-28^ In the cytosol, DELE1 oligomerizes and binds to the ISR kinase HRI, initiating HRI activation, eIF2α phosphorylation, and subsequent downstream ISR signaling.^25,26,29^ Thus, mitochondrial stress can activate ISR signaling through this OMA1-DELE1-HRI signaling axis. Apart from HRI, other ISR kinases including GCN2, PKR, and PERK can also be activated by mitochondrial insults, further highlighting a central role for the ISR in regulating mitochondria during conditions of stress.^30-32^

Other stress-responsive signaling pathways have also been proposed to regulate mitochondria in response to proteotoxic insult. The cytosolic heat shock response (HSR)-associated transcription factor HSF1 regulates expression of mitochondrial chaperones such as HSP60, suggesting an important role for HSF1 in adapting mitochondria proteostasis in response to stress.^33,34^ Consistent with this, HSF1 is activated when mitochondrial protein import decreases and mitochondrial ROS production increases.^35^ HSF1 was also previously suggested to be required for the induction of mitochondrial chaperones in response to mitochondrial proteotoxic stress.^34^ Disruptions of mitochondrial proteostasis can also lead to the activation of the oxidative stress response (OSR)-associated transcription factor NRF2. For example, genetic depletion of the mitochondrial chaperone HSP60 increases ROS and promotes NRF2 activation in clear cell renal cell carcinoma.^36^ Further, dysregulation of biological pathways localized to mitochondria such as the tricarboxylic acid (TCA) cycle lead to the accumulation of reactive metabolites that can activate NRF2 signaling.^37-40^ These results suggest that disruptions in mitochondrial proteostasis and function could induce activation of multiple different stress-responsive signaling pathways.

Here, we define the stress-responsive signaling pathways activated in response to mitochondrial proteostasis disruption induced by genetic perturbation of mitochondrial proteostasis. Towards that aim, we adapted and implemented a gene set profiling approach that monitors expression of gene sets comprised of transcriptional targets regulated downstream of stress-responsive signaling pathways including the ISR, HSR, OSR, and the endoplasmic reticulum (ER) unfolded protein response (UPR) in published CRISRPi Perturb-seq datasets from K562 cells.^41,42^ We initially verified this approach by monitoring the activation of our gene sets in cells where negative regulators of these pathways were depleted, confirming the ability for our gene sets to efficiently report on the activation of these pathways. We further validated our gene set approach by demonstrating that genetic depletion of ER proteostasis factors leads to preferential activation of the UPR, as predicted^43^, confirming our ability to identify stress-responsive signaling pathways preferentially activated by organelle-selective proteostasis disruptions. We then used this profiling strategy to monitor activation of stress pathway gene sets in cells CRISPRi-depleted of known mitochondrial proteostasis factors.^1^ Intriguingly, this identified the ISR as the sole stress-responsive signaling pathway activated by mitochondrial proteostasis disruption, with no other pathway showing significant activation. This selectivity was confirmed transcriptome-wide, indicating that mitochondrial proteostasis disruptions preferentially activate ISR signaling. We then expanded this study to show that CRISPRi-depletion of all mitochondrial proteins^44^ similarly demonstrates preferential activation of the ISR over other stress-responsive signaling pathways. Collectively, these results identify the ISR as a predominant stress-responsive signaling pathway activated in response to mitochondrial proteostasis stress, further motivating efforts to better understand the critical importance of this pathway in adapting mitochondrial proteostasis and function in the context of health and disease.

## RESULTS

### Transcriptional gene sets report on stress pathway activation in Perturb-seq datasets

We previously defined sets of 10-20 genes whose expression is regulated downstream of stress-responsive signaling pathways including the unfolded protein response (UPR; specifically the ATF6 and IRE1 arms of the UPR), integrated stress response (ISR), heat shock response (HSR), and oxidative stress response (OSR)(**Table S1**).^41^ We also defined a set of 19 control genes with expression that was relatively stable across diverse cellular insults (**Table S1**). Our stress pathway gene sets were largely defined using chemical genetic and genetic approaches that selectively activate stress-responsive signaling pathways independent of cellular stress to accurately monitor pathway activation in multiple model systems.^41^ Consistent with this, our gene sets have been used to report on the activation of stress-responsive signaling pathways in diverse cellular and in vivo models.^41,45-51^ However, the potential for these gene sets to report on the activity of stress-responsive signaling pathways in large scale genetic screens has not been previously explored.

Here, we defined the ability of our gene sets to report on activation of the UPR, ISR, HSR, and OSR in published Perturb-seq dataset where single cell RNAseq was performed on K562 cells CRISPRi-depleted of 11,258 individual genes.^42^ Initially, we confirmed that the expression of genes included in our stress pathway gene sets were not significantly altered in cells expressing non-targeting guide RNAs (**Fig. S1A-D**). Next, we monitored activation of our gene sets in K562 cells CRISPRi-depleted of genes encoding proteins involved in the regulation of these pathways. We initially focused on monitoring ISR activation in cells CRISPRi-depleted of genes that regulate the ISR. We monitored the expression of our ISR target gene set in cells CRISPR-deleted of genes encoding the upstream ISR transcription factor *ATF4*, the four ISR kinases (i.e., *EIF2AK1/HRI, EIF2AK2/PKR, EIF2AK3/PERK*, and*EIF2AK4/GCN2*,), the target of ISR kinases *EIF2A*, subunits of the guanine exchange factor EIF2B (i.e., *EIF2B1-EIF2B5*), and the two eIF2α phosphatases *PPP1R15A* and *PPP1R15B*.^13,14^ Depletion of *ATF4* reduced expression of our ISR target gene set, but not gene sets regulated by other stress-responsive signaling pathways (**Fig. 1A,B**). This is consistent with our ISR target genes being predominantly regulated downstream of ATF4. Depletion of EIF2B subunits, most notably *EIF2B3-5*, increased expression of our ISR target gene set (**Fig. 1A,C-D, Fig. S1E-G**). The depletion of these EIF2B genes did not induce expression of HSR or OSR target gene sets, although we did observe modest increases in the expression of target genes regulated by the UPR. This indicates that depletion of EIF2B factors preferentially activates the ISR, consistent with its role as a regulator of ternary complex formation.^52,53^ Collectively, these results indicate that our ISR target gene set effectively reports on ISR activity in this Perturb-Seq dataset.

**Figure 1.**
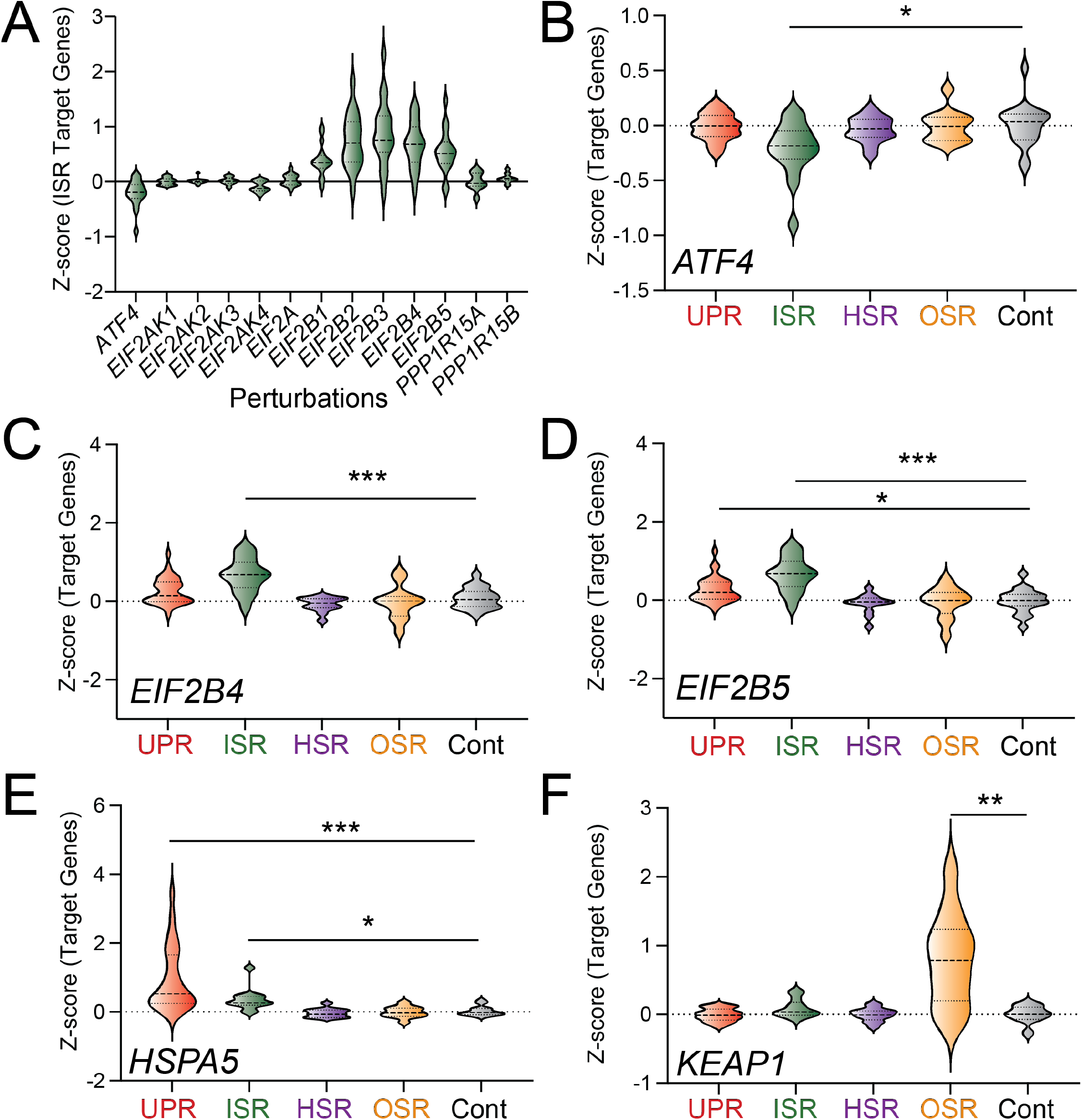
Target gene sets define stress pathway activation in Perturb-seq datasets. **A**. Expression, measured by z-score, of ISR target gene set comprising 13 genes in K562 cells CRISPRi-depleted of the indicated ISR component. The ISR target genes are shown in **Table S1. B-F**. Expression, measured by z-score, for UPR, ISR, HSR, OSR, and control target gene sets in K562 cells CRISPRi-depleted of *ATF4* (**B**), *EIF2B4* (**C**), *EIF2B5* (**D**), *HSPA5* (**E**), or *KEAP1* (**F**). The target gene sets used in this experiment were defined as in ^41^ and are shown in **Table S1**. *p<0.05, **p<0.01, ***p<0.005 for Brown-Forsythe and Welch ANOVA.

Next, we sought to confirm activation of other stress pathway gene sets by monitoring their expression in K562 cells CRISPRi-depleted of genes encoding proteins that negatively regulate these pathways. The ER HSP70 chaperone BiP/HSPA5 negatively regulates the three signaling arms of the UPR activated downstream of the ER stress sensors IRE1, ATF6, and PERK, the latter also being a component of the ISR.^54,55^ Previous results have shown that genetic depletion of *HSPA5* activates all three arms of the UPR.^43^ Thus, we predicted that CRISPRi-depletion of *HSPA5* should selectively activate the UPR (reflecting IRE1 and ATF6 activation) and the ISR (reflecting PERK activation). As expected, CRISPRi-depletion of *HSPA5* increased expression of the UPR and ISR gene sets (**Fig. 1E**). The E3 ubiquitin ligase KEAP1 negatively regulates NRF2-dependent transcriptional regulation of the OSR.^56,57^ Thus, we predicted that CRISPRi-depletion of *KEAP1* should activate the NRF2-regulated OSR. As expected, CRISPRi-depletion of *KEAP1* selectively increased expression of the OSR target gene set (**Fig. 1F**). Further, CRISPRi-depletion of *UBA1* – an E1 ubiquitin ligase involved in the ubiquitin-proteasome pathway – increased expression of HSR target genes (**Fig. S1H**), reflecting the HSR activation induced by reduced proteasomal degradation.^58^ Collectively, these results indicate that our gene sets efficiently report on the activation of these stress-responsive signaling pathways in Perturb-seq datasets from CRISPRi-depleted K562 cells.

### Genetic depletion of ER proteostasis factors selectively activates UPR transcriptional signaling

We next wanted to validate the ability for our gene sets to identify stress-responsive signaling pathways induced by perturbations to the proteostasis environment of a specific organelle. Genetic disruption of ER proteostasis selectively activates the UPR.^43^ Thus, we wanted to demonstrate that our gene sets similarly show selective UPR activation induced by ER proteostasis perturbations. We monitored the expression of stress pathway target genes in K562 cells CRISPRi-depleted of the 215 enumerated proteins comprising discrete ER proteostasis pathways including ER chaperones/folding enzymes, glycoproteostasis factors, protein degradation pathways, and protein targeting/trafficking pathways.^1^ We confirmed the knockdown efficiency for these ER proteostasis factors (**Fig. S2**). Monitoring the expression of individual target genes shows that UPR target genes are preferentially induced by CRISPRi-depletion of ER proteostasis factors (**Fig. 2A**). To further define pathway activation, we ascribed a stress pathway activation score, defined by the average z-score for target genes comprising an individual stress pathway gene set, for cells CRISPRi-depleted of individual ER proteostasis factors. This showed that perturbations to multiple different ER proteostasis factors including the ER HSP70 chaperone *HSPA5* (see also **Fig. 1E**), the oligosacharyl transferase subunits *DDOST1* and *DAD1*, and the ER protein targeting factors *SRP68* and *SRP72* activated the UPR target gene set (**Fig. 2B**). We also observed modest activation of the ISR target gene set in cells CRISPRi-depleted of these ER proteostasis factors, reflecting activation of the PERK signaling arm of the UPR/ISR. However, no other stress pathways were activated in cells depleted of ER proteostasis factors. This indicates that, as predicted, organelle-specific disruption of ER proteostasis leads to preferential activation of the UPR and can be reliably detected by our target gene sets.

**Figure 2.**
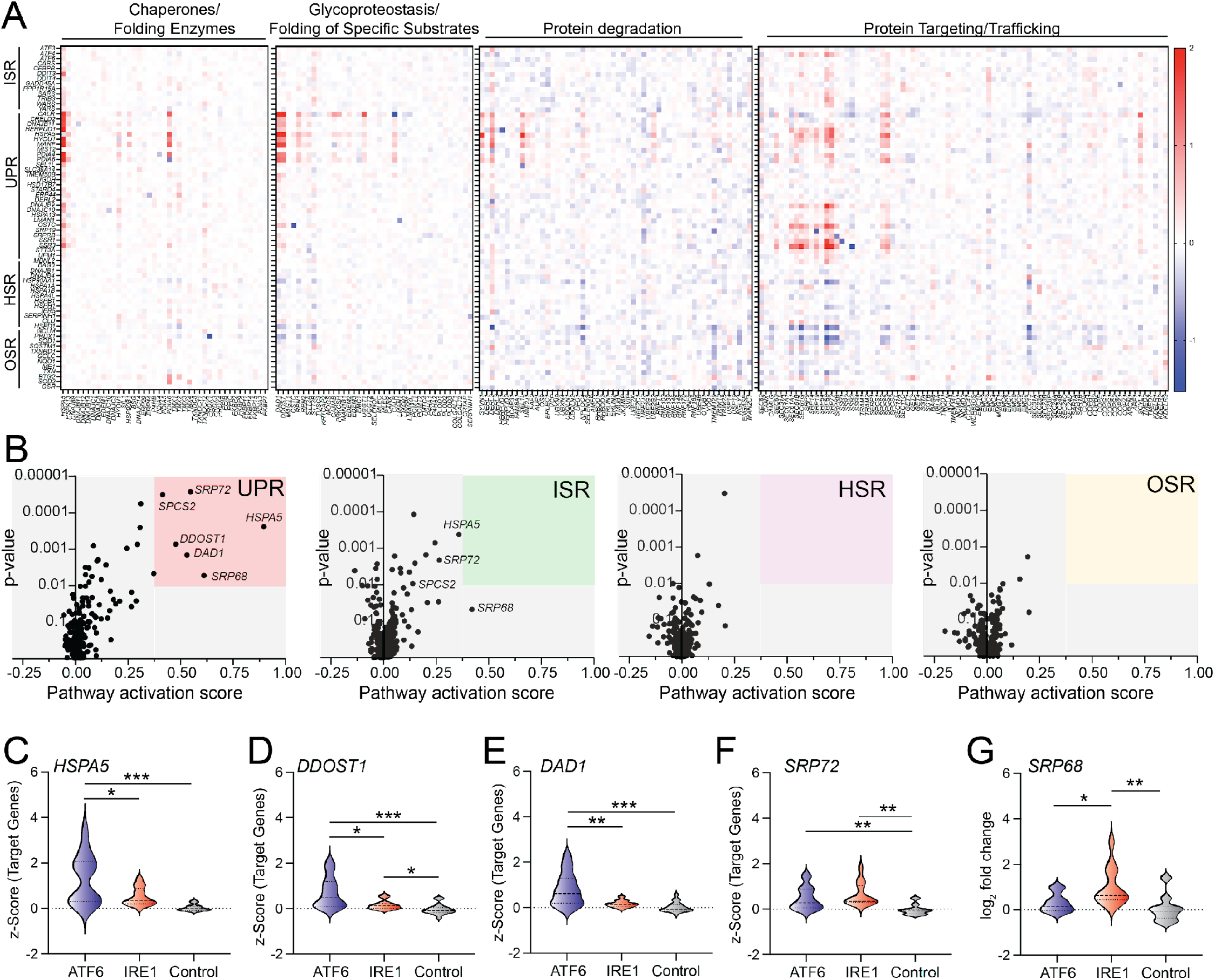
ER proteostasis perturbations selectively activate the unfolded protein response. **A**. Heat map showing the expression, measured by z-score, of ISR, UPR, HSR, and OSR target genes in K562 cells CRISPRi-depleted of components of the indicated ER proteostasis pathway, as defined in ^1^. **B**. Pathway activation score vs p-value for UPR, ISR, HSR, and OSR target genes in K562 cells CRISPRi-depleted of individual ER proteostasis factors. Pathway activation score was defined by the average z-score for pathway-specific target genes and p-values were calculated by comparing the expression of pathway specific target gene set to the expression of the control gene sets. **C-G**. Expression, measured by z-score, for ATF6, IRE1/XBP1s, and control target gene sets in K562 cells CRISPRi-depleted of *HSPA5* (**C**), *DDOST1* (**D**), *DAD1* (**E**), *SPR72* (**F**), or *SRP68* (**G**). ATF6 and IRE1/XBP1s gene sets are defined in **Table S1**. *p<0.05, **p<0.01, ***p<0.005 for Brown-Forsythe and Welch ANOVA.

The UPR gene set comprises target genes regulated downstream of both the IRE1/XBP1s and ATF6 arms of the UPR.^41^ Intriguingly, perturbations to specific ER proteostasis factors appear to differentially impact the expression of individual UPR target genes (**Fig. 2A**). This suggests that these perturbations may be differentially influence activation of ATF6 and IRE1/XBP1s signaling. To further probe this, we monitored the expression of UPR target genes predominantly regulated by ATF6 or IRE1/XBP1s signaling, as previously defined^41^, to evaluate the differential regulation of these two UPR pathways induced by specific perturbations. Depletion of *HSPA5, DAD1*, and *DDOST1* preferentially increased expression of target genes regulated downstream of ATF6, as compared to those regulated downstream of IRE1 (**Fig. 2C-E**). In contrast, depletion of *SRP68* and *SPR72* – two genes involved in the co-translational targeting of proteins to the ER^59,60^ – result in preferential activation of IRE1 target genes (**Fig. 2F,G**). This is consistent with previous reports showing that disruption of ER protein targeting preferentially induces IRE1 activation.^43^ These results demonstrate the potential for our gene set approach to define differential activation of UPR signaling arms induced by distinct perturbations. Moreover, these results establish the ability for our gene sets to identify stress-responsive signaling pathways activated by perturbations of proteostasis within a specific organelle using Perturb-seq data.

### Genetic depletion of mitochondrial proteostasis factors preferentially activates the ISR

Previous results have suggested that disruptions in aspects of mitochondrial proteostasis leads to the activation of multiple stress-responsive signaling pathways including the ISR, HSR, and OSR.^25-28,30-32,34-40^ Here, we sought to probe the activation of these stress-responsive signaling pathways in K562 cells CRISPRi-depleted of individual mitochondrial proteostasis factors. The mitochondrial proteostasis network comprises 94 chaperones/folding enzymes, proteases, and import factors^1^, 82 of which were included in the published Perturb-seq dataset.^42^ We confirmed efficient knockdown efficiency of these genes (**Fig. S3A**).

We monitored the expression of stress pathway target genes in cells depleted of these mitochondrial proteostasis factors. Genetic reductions in numerous mitochondrial proteostasis factors including the mitochondrial chaperones *HSPA9* and *HSPE1* and the mitochondrial protein import proteins *TIMM23B* and *TOM22* increased activation of the ISR target genes (**Fig. 3A-C**). No other stress pathway gene set was activated upon depletion of mitochondrial proteostasis factors (**Fig. 3A-C**). The activation of the ISR target gene set appeared largely independent of knockdown efficiency, as genes with relatively low knockdown efficiency (e.g., *TIMM23B, PHB2*) still showed activation of this pathway (**Fig. S3B**). These results indicate that the ISR target genes are preferentially induced by CRISPRi-depletion of mitochondrial proteostasis factors in these cells.

**Figure 3.**
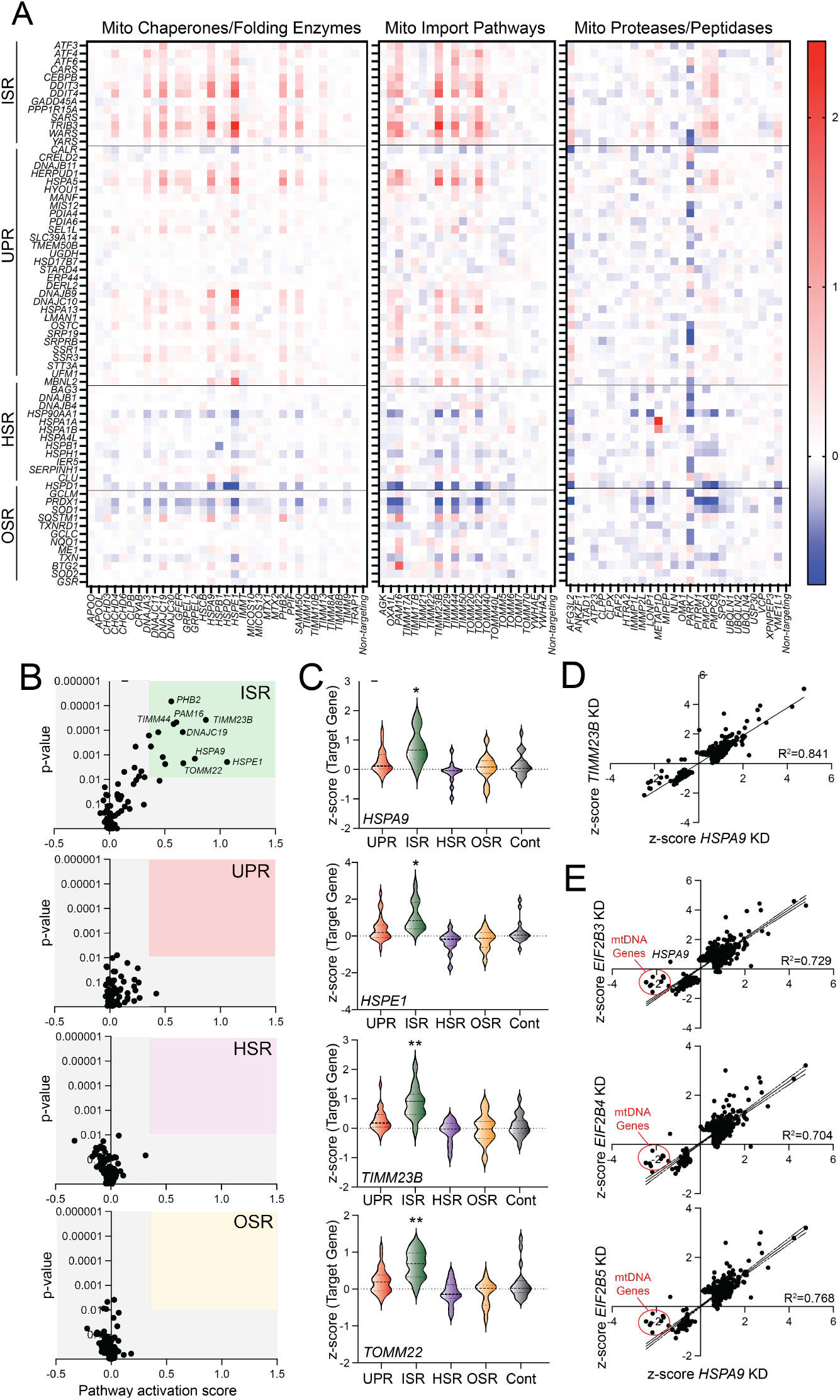
Mitochondrial proteostasis gene perturbation preferentially activates the ISR. **A**. Heat map showing the expression, measured by z-score, of ISR, UPR, HSR, and OSR target genes in K562 cells CRISPRi-depleted of components of the indicated mitochondrial proteostasis pathway, as defined in ^1^. **B**. Pathway activation score vs -log p-value for UPR, ISR, HSR, and OSR target genes in K562 cells CRISPRi-depleted of individual mitochondrial proteostasis factors. Pathway activation score was defined by the average z-score for pathway-specific target genes. P-values were calculated by comparing the pathway specific target gene set expression to the expression of the control gene sets. **C**. Expression, measured by z-score, for UPR, ISR, HSR, OSR, and control target genesets in K562 cells CRISPRi depleted of *HSPA9, HSPE1, TIMM23B*, or *TOMM22*, as indicated. *p<0.05, **p<0.01, ***p<0.005 for Brown-Forsythe and Welch ANOVA, as compared to the control geneset. **D**. Comparison of gene expression, as measured by z-score, in K562 cells CRISPRi-depleted of either *HSPA9* or *TIMM23B*. Only genes increased or decreased |value| > 0.5 in response to either perturbation are shown. **E**. Comparison of gene expression, measured by z-score, in K562 cells CRISPRi-depleted of *HSPA9* or *EIF2B3, EIF2B4, or EIF2B5*, as indicated. Only genes increased or decreased |value| > 0.5 in response to either perturbation are shown.

To further probe the transcriptome-wide impact of mitochondrial proteostasis factor depletion, we compared the expression of genes with z-scores >0.5 or <0.5 in K562 cells CRISPRi depleted of *HSPA9, HSPE1*, or *TIMM23B*. Intriguingly, all these perturbations induced similar transcriptome-wide changes in gene expression, reflected by the linear relationship between gene expression changes observed in cells depleted of these individual mitochondrial proteostasis factors (**Fig. 3D, Fig. S3C-D**). GO analysis also demonstrates that these perturbations increase and decrease expression of genes involved in similar biological pathways related to the ISR such as response of EIF2AK4 to amino acid deficiency, cellular response to starvation, and the unfolded protein response (**Fig. S3E**). We next compared the gene expression changes in cells CRISPRi-depleted of *HSPA9* or components of the eIF2B complex – the latter of which preferentially activate the ISR (**Fig. 1A**). This demonstrated that these two perturbations, which induce ISR signaling through distinct mechanisms, showed similar gene expression changes (**Fig. 3E**). This is in contrast to the distinct transcriptional profiles observed when comparing gene expression changes in cells CRISPRi-depleted of *HSPA9* and *UBA1* (**Fig. S3F**). We did note that expression of genes encoded by mitochondrial DNA were selectively reduced in *HSPA9*-depleted cells, as compared to cells depleted of EIF2B subunits, potentially reflecting a disruption in mitochondrial content and/or activity selectively induced by mitochondrial proteostasis disruption (**Fig. 3E**). Regardless, our results indicate that disruptions in mitochondrial proteostasis afforded by perturbations such as *HSPA9* depletion demonstrate transcriptome-wide selectivity for ISR activation that can be mimicked by non-mitochondrial perturbations to other biological pathways that preferentially induce the ISR.

### Depletion of other mitochondrial proteins preferentially activates the ISR

We next sought to expand this approach to probe the expression of stress pathway gene sets in cells depleted of other types of mitochondrial proteins. We collated the expression profiles for K562 cells CRISPRi-depleted of all mitochondrial proteins, as defined by MitoCarta 3.0.^44^ Principal component analysis (PCA) suggested that there was minimal variation (<13%) across all perturbations to mitochondrial proteins and no discrete clustering (**Fig. S4A**). To determine the extent to which ISR activation corresponded with the transcriptional profiles of mitochondrial perturbations, we also included EIF2B gene perturbations, which promote ISR activity through a distinct mechanism. The PCA suggests that the extent of ISR activation is associated with the transcriptomic variability observed across all mitochondrial perturbations. (**Fig. S4A**). Consistent with this, using our gene sets to monitor activation of different stress responsive signaling pathways, we found only the ISR showed significant activation in response to mitochondrial gene perturbation (**Fig. 4A**). Unsurprisingly, the majority of genes whose perturbation led to ISR activation were mitochondrial proteostasis factors, reflecting the selectivity for ISR activation to disruption in mitochondrial proteostasis. However, we also identified other mitochondrial proteins whose depletion activated the ISR. These include the tRNA synthetase *IARS2*, the mitochondrial phospholipid transporter *PRELID3B*, and the mitochondrial coA transporter *SLC25A42* (**Fig. 4B**). GO analysis of gene expression changes induced by perturbation to *IARS2, PRELID3B*, and *SLC25A42* showed preferential activation of ISR-associated terms (**Fig. S3E**). This indicates that these perturbations appear to preferentially induce ISR signaling. Further, comparing the gene expression changes induced by depletion of *IARS2, PRELID3B*, or *SLC25A42* to those induced by *EIF2B5* depletion demonstrated strong correlations, indicating that all these perturbations induce similar gene expression changes (**Fig. 4C** and **Fig. S4B,C**). These results further identify the ISR as the predominant stress responsive signaling pathway activated in response to disruptions in mitochondrial proteostasis and function.

**Figure 4.**
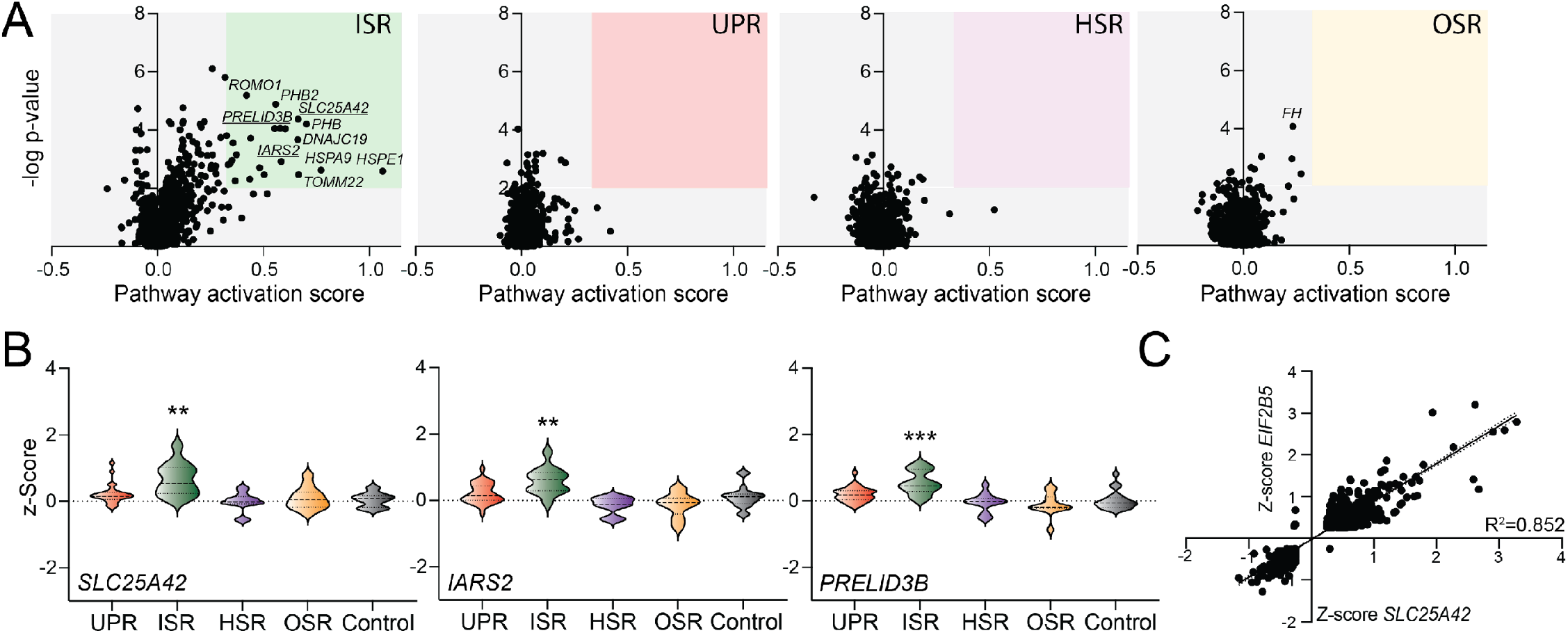
Mitochondrial gene perturbations preferentially activate the ISR. **A**. Pathway activation score vs -log p-value for UPR, ISR, HSR, and OSR target genes in K562 cells CRISPRi-depleted of genes encoding mitochondrial proteins, as defined in ^44^. Pathway activation score was defined by the average z-score for pathway-specific target genes. P-values were calculated by comparing the pathway specific target gene expression to the expression of the control gene sets. **B**. Expression, measured by z-score, for UPR, ISR, HSR, OSR, and control target gene sets in K562 cells CRISPRi depleted of *SLC25A42, IARS2*, or *PRELID3B*, as indicated. **p<0.01, ***p<0.005 for Brown-Forsythe and Welch ANOVA, as compared to the control geneset. **C**. Comparison of gene expression, as measured by z-score, in K562 cells CRISPRi-depleted of either *SLC25A42* or *EIF2B5*. Only genes increased or decreased |value| > 0.5 in response to either perturbation are shown.

## DISCUSSION

Here, we adapt and implement a gene set profiling approach to probe the activation of stress-responsive signaling pathways in K562 cells where organellar proteostasis is disrupted by CRISRPi-depletion of proteostasis factors. We establish the validity of this approach by demonstrating the ability of our gene sets to accurately report on stress pathway activation and confirming the sensitivity of UPR activation to ER proteostasis disruption. Further, we show that genetic perturbations to mitochondrial proteostasis and function leads to preferential activation of the ISR target gene set, with no other pathway showing activation. These results identify the ISR as the predominant stress-responsive signaling pathway activated in response to genetic depletion of mitochondrial proteostasis factors.

While mitochondrial perturbations can activate all four ISR kinases, recent results have coalesced around a model whereby cells primarily respond to mitochondrial stress through the activation of the OMA1-DELE1-HRI ISR signaling axis.^25-29^ The elucidation of this pathway has revealed a clear molecular mechanism by which cells communicate mitochondrial perturbations to the nucleus and initiate adaptive mitochondrial remodeling through the ISR. Consistent with this, ISR activation has been shown to regulate diverse aspects of mitochondrial biology including proteostasis, morphology, energy metabolism, lipid metabolism, and apoptotic signaling.^15-24^ Thus, combined recent results demonstrating both a signaling mechanism to activate the ISR in response to mitochondrial stress and an adaptive program to promote mitochondrial remodeling following pathologic insults highlights a central role for the ISR in regulating mitochondria in response to stress. Our work further highlights the central importance of the ISR for mitochondrial regulation, identifying the ISR as the only stress responsive signaling pathway activated by perturbations to mitochondrial proteostasis and function.

Surprisingly, we did not observe the activation of other stress-responsive signaling pathways in cells depleted of mitochondria proteostasis factors. Notably, we did not see activation of the HSR, which has been previously suggested to be activated in response to perturbations to mitochondrial proteostasis.^34,35^ One potential explanation for this observation is that the HSR is only activated by acute disruptions of mitochondrial proteostasis, while more chronic disruptions, such as those resulting from CRISPRi depletion, promote adaptive cellular remodeling that both reduces the burden associated with mitochondrial proteotoxic stress. Thus, in this model, HSR activation would only be observed in cells subjected to acute mitochondrial proteostasis stress such as that afforded by pharmacologic inhibition of mitochondrial proteostasis factors. Consistent with this, many of the experiments showing HSR activation induced by mitochondria stress utilize small molecule inhibitors of mitochondrial proteostasis factors to induce acute proteotoxic stress (e.g., GMPP to inhibit the mitochondrial chaperone TRAP1).^34,35^ Alternatively, the lack of other stress pathways being activated by mitochondrial proteostasis disruption could reflect cell type differences in the response to mitochondrial stress. Regardless, the selectivity for ISR activation in this study highlights the central importance of this pathway in regulating mitochondria in response to chronic stress.

Our identification of the ISR as the predominant stress-responsive signaling pathway activated by mitochondrial proteotoxic stress underscores a unique opportunity to target the ISR to correct pathologic mitochondrial dysfunction in etiologically diverse diseases. For example, recent work showed that pharmacologic activation of the ISR kinase GCN2 reduced mitochondrial fragmentation and respiratory chain dysfunction in cells deficient in the ISR kinase PERK^18^ – a condition linked to many neurodegenerative diseases including Alzheimer’s disease and Progressive Supranuclear Palsy.^61,62^ Further, the potential to pharmacologically access adaptive mitochondrial remodeling induced by ISR activation could allow correction of mitochondrial dysfunction induced by genetic, environmental, or aging-related insults. Thus, as we, and others, continue probing the activation and activity of the ISR in the context of mitochondrial stress, we anticipate identifying many additional opportunities to correct pathologic mitochondrial dysfunction implicated in many other types of diseases.

## METHODS

### Data acquisition and code availability

Processed pseudobulk perturb-seq datasets for K562 and were obtained from the gwps.wi.mit.edu figshare data portal. The K562 dataset comprises sequencing data from 8,248 mRNA transcripts obtained 8 days after transduction for 11,258 sgRNAs (“pertubations”). The pseudobulk data files report the z-normalized expression (“z-score”) as the standard deviation of each gene relative to the mean and standard deviation of a control set of sgRNAs from the same single cell library preparation, or 10X gemgroup. Data files were downloaded as an .h5ad file read using SCANPY (version 1.6) and AnnData (version 0.10). Knockdown efficiencies, as indicated in figures, were stored in the pseudobulk .h5ad file as ‘fraction knockdown’ in the obs dataframe. Code for subsequent analysis steps are available upon reasonable request. A comprehensive list of ER and mitochondrial proteostasis factors, obtained from the Proteostasis Network and Mitocarta 3.0, was used to filter data for the specific perturbations indicated for the graphs shown.

### Gene ontology (GO) enrichment analysis

The GO analysis in Fig. S3E for multiple perturbations was performed and visualized using ClusterProfiler (version 4.10.0) and the reactome database for all transcripts with an |value| > 1. The top 5 enriched GO terms were visualized.

### Principal Component Analysis

The *prcomp* command from the R stats package (version 4.3.2) was used to perform principal component analysis (PCA) and the factoextra package (version 1.0.7) was used for visualization. PCA was performed using all quantified transcripts detected in the pseudobulk data of all cells expressing sgRNAs targeting mitochondrial proteins listed in Mitocarta 3.0 or sgRNAs targeting EIF2B subunits.

### Statistics

All statistical analyses were performed using Prism 9 (GraphPad, San Diego, CA) as described. Simple linear regressions were used to generate the R2 values and define the correlations for two sgRNA datasets for all transcripts with |value| > 0.5. Brown-Forsythe and Welch ANOVA statistical tests were used to determine statistically different SDs and detect statistically significant changes in stress signaling pathway activity, respectively, s= with post hoc testing to define specific statistical relationships.

## ACKNOWLEDGEMENTS

We thank Sergei Kutseikin for critical reading of this manuscript. This work was funded by the National Institutes of Health (NS095892, NS125674 to RLW).

---

**Figure S1 (Supplement to Figure 1).**
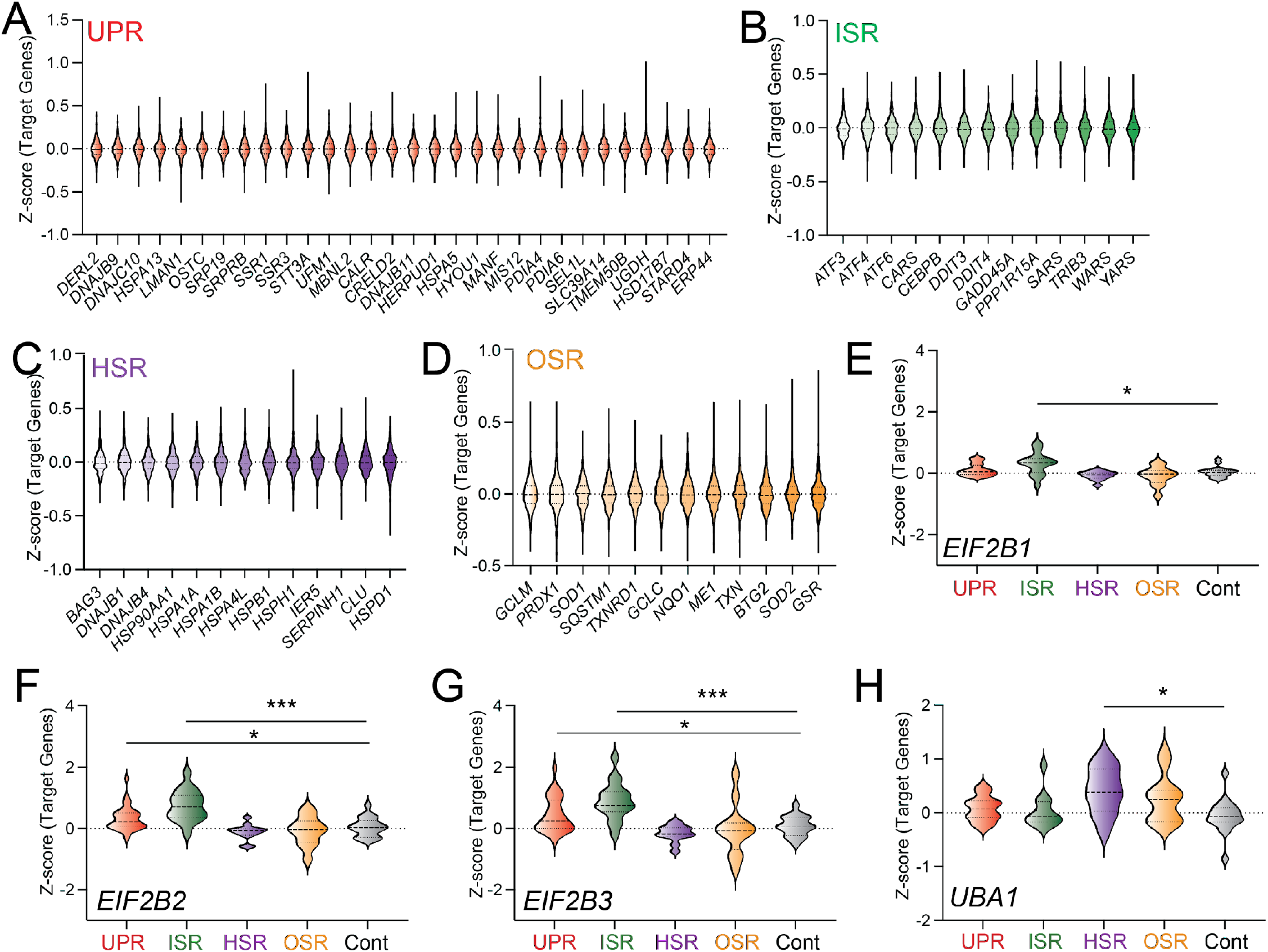
**A-D**. Expression, measured by z-score, of UPR (**A**), ISR (**B**), HSR (**C**), and OSR (**D**) in K562 cells expressing non-silencing guide RNAs. **E-H**. Expression, measured by z-score, of UPR, ISR, HSR, OSR, and control gene sets in K562 cells CRISPRi-depleted of *EIF2B1* (**E**), *EIF2B2* (**F**), *EIF2B3* (**G**), or *UBA1* (**H**). *p<0.05, ***p<0.005 for Brown-Forsythe and Welch ANOVA.

**Figure S2 (Supplement to Figure 2).**
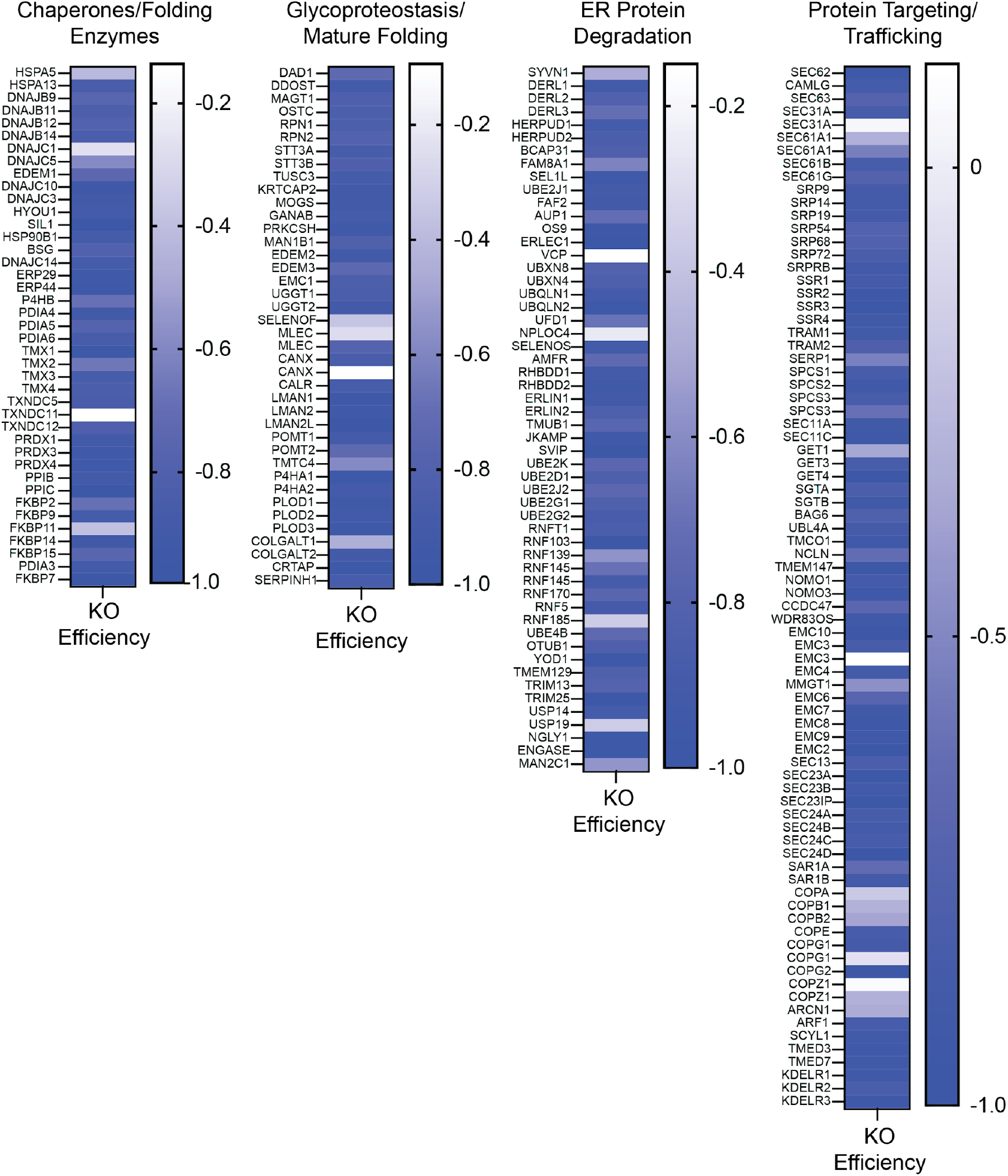
Knockdown efficiency of ER proteostasis genes. ER proteostasis genes were defined as in ^1^. Knockdown efficiency was defined as in ^42^.

**Figure S3 (Supplement to Figure 3).**
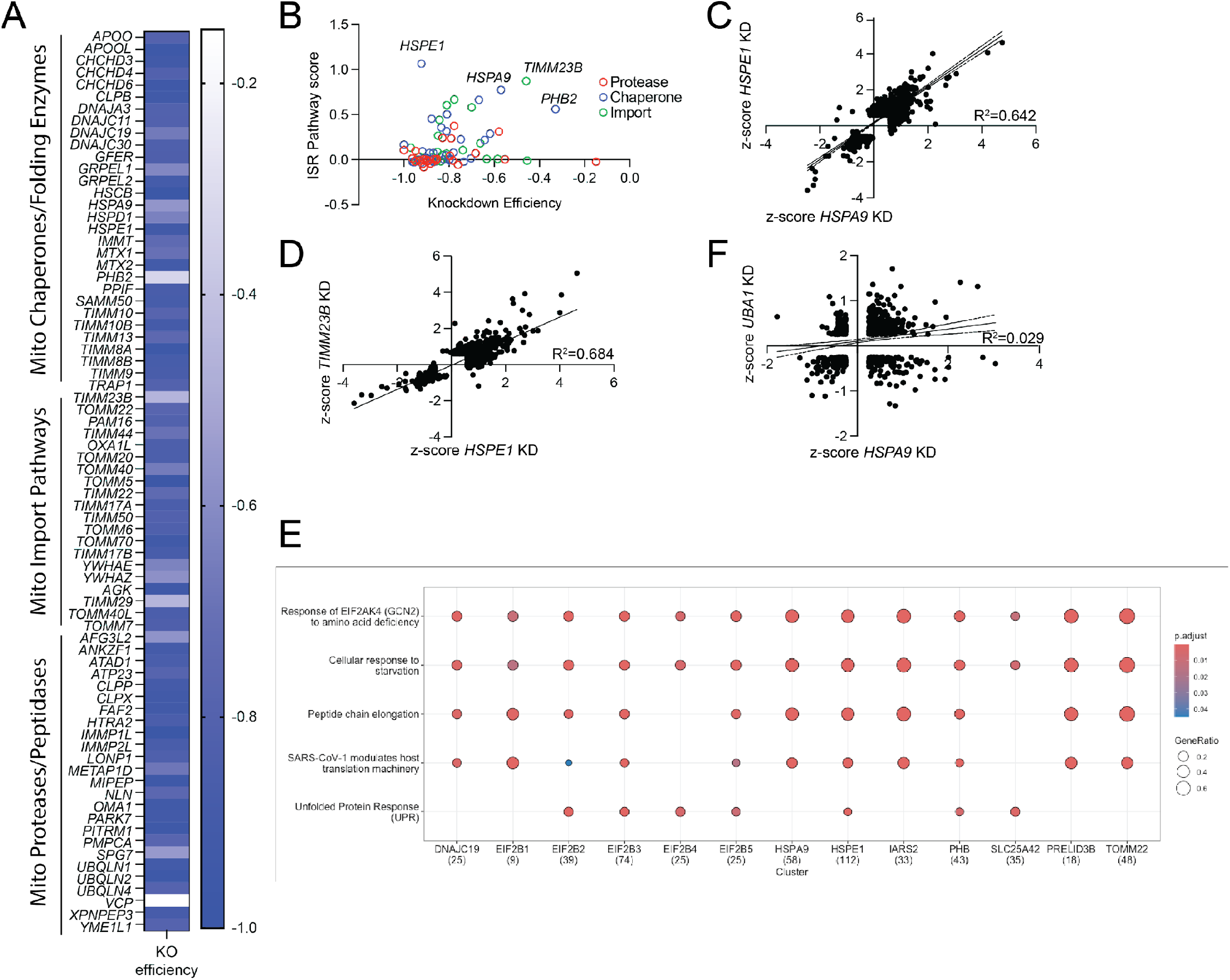
**A**. Knockdown efficiency of ER proteostasis genes. ER proteostasis genes were defined as in ^1^. Knockdown efficiency was defined as in ^42^. **B**. Plot of knockdown efficiency vs. ISR pathway score for mitochondrial proteostasis factors. IS activation score was defined by the average z-score for ISR target genes. **C**,**D**. Comparison of gene expression, as measured by z-score, in K562 cells CRISPRi-depleted of *HSPA9* vs *HSPE1* (**C**) or *TIMM23B* (**D**). Only genes increased or decreased |value| > 0.5 in response to either perturbation are shown. **E**. Comparison of ISR-related GO terms induced in K562 cells CRISPRi-depleted of the indicated mitochondrial gene. **F**. Comparison of gene expression, as measured by z-score, in K562 cells CRISPRi-depleted of *HSPA9* vs *UBA1*. Only genes increased or decreased |value| > 0.5 in response to either perturbation are shown.

**Figure S4 (Supplement to Figure 4).**
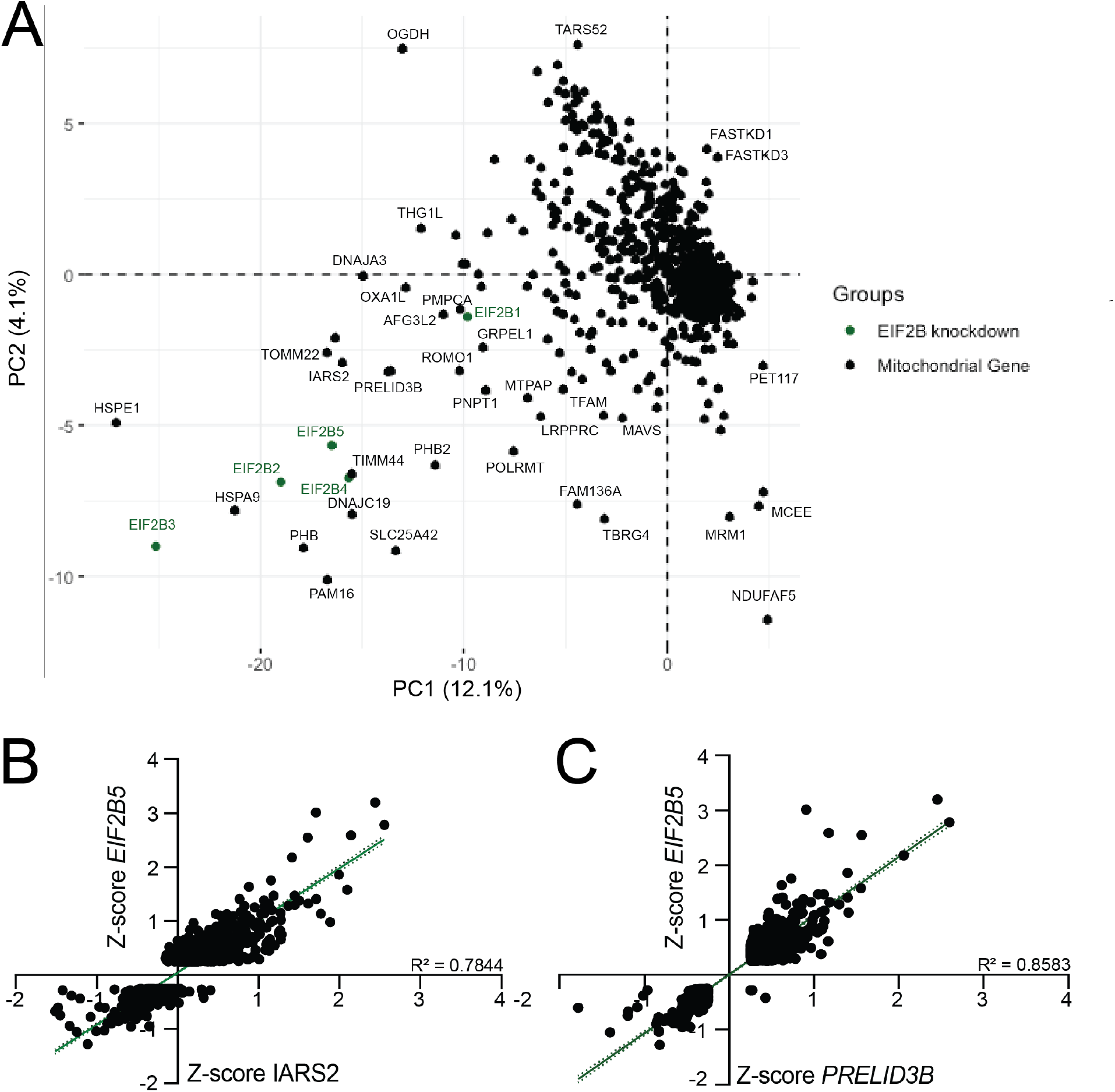
**A**. Principal component analysis (PCA) for gene expression observed in K562 cells CRISPRi-depleted of individual genes encoding mitochondrial proteins. Components of the EIF2B guanine exchange factor (*EIF2B1-5*) are also shown in green. **B**,**C**. Comparison of gene expression, as measured by z-score, in K562 cells CRISPRi-depleted of *IARS2* (**B**) or *PRELID3B* (**C**) vs *EIF2B*. Only genes increased or decreased |value| > 0.5 in response to either perturbation are shown.

